# Druggability Assessment in TRAPP using Machine Learning Approaches

**DOI:** 10.1101/2019.12.19.882340

**Authors:** Jui-Hung Yuan, Sungho Bosco Han, Stefan Richter, Rebecca C. Wade, Daria B. Kokh

## Abstract

Accurate protein druggability predictions are important for the selection of drug targets in the early stages of drug discovery. Due to the flexible nature of proteins, the druggability of a binding pocket may vary due to conformational changes. We have therefore developed two statistical models, a logistic regression model (TRAPP-LR) and a convolutional neural network model (TRAPP-CNN), for predicting druggability and how it varies with changes in the spatial and physicochemical properties of a binding pocket. These models are integrated into TRAPP (TRAnsient Pockets in Proteins), a tool for the analysis of binding pocket variations along a protein motion trajectory. The models, which were trained on publicly available and self-augmented data sets, show equivalent or superior performance to existing methods on test sets of protein crystal structures, and have sufficient sensitivity to identify potentially druggable protein conformations in trajectories from molecular dynamics simulations. Visualization of the evidence for the decisions of the models in TRAPP facilitates identification of the factors affecting the druggability of protein binding pockets.

## Introduction

Drug development is a very costly and time-consuming process, with only 7-11% of compounds in phase I testing finally becoming approved drugs.^1^ A thorough drug target validation in the earlier stages of drug discovery projects could reduce the waste of resources and improve the success rate. Thus, the druggability of a protein,^2–4^ which describes the protein’s potential to accommodate low molecular weight drug molecules that modulate its biological function, should be evaluated during target validation. Low molecular weight compounds may show a therapeutic effect when they bind with high affinity to a druggable disease-related protein at a specific position, known as the binding pocket or binding cavity. Information on protein druggability is particularly useful for prioritizing therapeutic targets, and for guiding the design of new drugs to avoid side-effects or unwanted polypharmacology.

The concept of druggability is used to described a biological target that is disease-related and can be modulated by commercially viable compounds that are usually orally available and are known as drug-like molecules.^5,6^ An empirical rule for evaluating the druglikeness of a given compound is Lipinski’s rule-of-five (RO5),^7^ which was derived based on the physicochemical properties of 90% of orally active drugs that achieved Phase II status. Thus, a more precise and practical definition of protein druggability can be formulated as the protein’s ability to bind drug-like ligands with high affinity. Most experimental approaches for assessing druggability have involved high-throughput or NMR-based screening using libraries of molecules that conform to lead-like characteristics.^3,8^ However, the high cost of instrumentation and the uncertainty of negative assessments due to potentially inadequate compound collections restrict the scope of experimental approaches. Therefore, computational procedures, typically based on virtual screening or machine learning,^9^ have been developed to provide a more generally applicable approach.

Here, we focus on protein structure-based druggability prediction using machine learning approaches to take advantage of the large amount of structural data on proteins. Existent statistical models for druggability prediction typically have 3 essential elements: an informative featurization of protein binding pockets, a druggability dataset, and a machine learning approach. The featurization usually involves two steps. First, a cavity or crevice in the protein is defined using a pocket estimation method. Then, the features are derived for the specific cavity. A wide range of features or descriptors can be used to characterize a pocket, including geometric information such as pocket volume or surface area, and physicochemical properties like hydrophobicity or polarity. The features that enable better discrimination between druggable and less-druggable pockets are then used as the input for the druggability prediction model. Based on the knowledge contained in the druggability dataset, the model learns to capture the relation between input features and output druggability using machine learning approaches. In order to compare the performance of different methods, consensus datasets, such as the non-redundant druggable and less druggable (NRDLD) dataset,^10^ have been developed.

One of the earliest methods, developed by Hajduk et al., ^3^ predicts experimental screening hit rate, as a druggability measure, with a linear regression model based on descriptors of the polar/apolar surface area, the surface complexity, and the pocket dimensions. For DrugPred,^10^ partial least-squares projection to latent structures discriminant analysis (PLSDA) was used to select 5 out of 22 descriptors that show significant contributions to the model decision with positive contributions to druggability from hydrophobicity and protein-ligand contact surface and a negative contribution from polar surface area. Volsite ^6^ encodes both the shape and the pharmacophoric properties of a binding cavity on regularly spaced grid points and derives 73 global and local descriptors from the grids, using a support vector machine algorithm (SVM) with a nonlinear kernel to train the druggability prediction model. The non-linearity of the SVM means that the contribution of each feature to the model cannot be obtained directly. Another grid-based method is DoGSiteScorer, ^11,12^ which maps the protein of interest onto a grid and estimates pockets and subpockets using a Difference of Gaussian filter.^13^ Based on the pocket-forming grid points and the pocket-lining residues, global and local descriptors are computed. From 17 global descriptors three descriptors namely, pocket depth, number of apolar amino acid residues and the pocket volume, are used to build a SVM model with a Gaussian kernel. In addition, local pocket properties are described by distance dependent histograms between atom pairs, e.g. hydrophilic-hydrophilic and lipophilic-lipophilic atom pairs, which allows druggability assessment through a nearest neighbor search. The local descriptors reveal that druggable pockets tend to have fewer short-range hydrophilic interaction pairs and more short-range lipophilic pairs. The combination of global and local predictors increases the reliability of the model. Subsequently, as the druggability prediction of a single target varies when applying different pocket estimation methods, PockDrug^14^ was developed to overcome these uncertainties by optimizing the model with different pocket estimation methods. It performs druggability prediction based on the consensus of seven linear discriminant analysis (LDA) models with complementary input features. In general, the properties involved describe the geometry, hydrophobicity and aromaticity of the pockets, as for most other druggability prediction models.^15,16^

Most druggability prediction methods have been developed to use the static structures of proteins determined by experimental methods, such as X-ray crystallography, NMR, or cryo-electron microscopy. However, proteins are dynamic and possess an inherent flexibility^17^ which alters the shape and properties of their binding pockets. ^18–20^ Therefore, it is valuable to explore the different conformations of a binding pocket by molecular simulation and the corresponding variations in druggability. Some methods that combine pocket druggability prediction and molecular dynamics simulation have been developed, such as MDpocket^21^ and JEDI. ^22^ To compute the druggability score of a complete trajectory efficiently, these methods use only a few descriptors, such as volume and hydrophobicity, and therefore might not be sensitive enough to capture the variations in druggability due to subtle conformational changes. Therefore, based on the assumption that a change in the conformation of a binding site leads to a change in its druggability, our aim in this work was to build druggability assessment models that are able to trace the fluctuation of druggability as a result of conformational changes. TRAPP is a tool that allows the exploration of different protein conformations, the analysis of binding pocket flexibility and dynamics, and the extraction of spatial and physicochemical information on the binding pocket conformations. The goal of this study was to make use of the information provided by TRAPP for assessing the druggability of a binding pocket as its shape changes using machine-learning based approaches. Global descriptors of the pocket were generated as the input for linear models using logistic regression (LR) and a support vector machine (SVM), whereas a grid representation of the pocket provided input to a convolutional neural network (CNN) for druggability prediction. Visualization tools were developed to elucidate the evidence for the decisions of the models. The performance of the models was evaluated and compared with other methods to predict druggability and they were found to perform equally well or better.

## Methods

### Data sets

Two data sets of experimentally determined protein structures were used to train and validate the druggability assessment models. In addition, to illustrate the capabilities of our models, MD trajectories of pocket motions were generated.

### NRDLD dataset

The Non-Redundant Druggable and Less Druggable (NRDLD) dataset is the largest publically accessible druggability dataset available to date.^10^ It contains 113 instances of proteins with high resolution (less than 2.6 Angstroms) crystal structures of which 71 are labeled as druggable and 42 are labeled as less-druggable. A protein is defined as druggable if it is able to noncovalently bind small drug-like ligands that are orally available and do not require administration as prodrugs, whereas a protein is less-druggable if none of the ligands with which it was co-crystallized simultaneously met the following three requirements: (1) satisfied Lipinski’s rule of five for orally available drugs; ^7^ (2) had *clogP* ≥ −2;^23^ and (3) The ligand efficiency was ≥ 0.3 kcal mol^−1^/heavy atom.^24^ The redundant instances were removed by choosing a single crystal structure to represent protein targets sharing a pairwise sequence identity of greater than 60%. The NRDLD data set has a pre-defined split, with a training set with 76 instances and a test set with 37 instances. Since the performance of the latest druggability prediction tools was mainly assessed using the NRDLD test set, it allows comparison of the performance of our method with other methods.

### DaPB dataset

A larger dataset, the Druggability augmented from PDBbind (DaPB) dataset, was constructed by filtering the 2017 release of the PDBbind refined set. ^25^ The PDBbind database collects the experimentally measured binding affinity data for the biomolecular complexes in the Protein Data Bank (PDB), and the PDBbind refined set is a subset containing 4154 protein-ligand complexes with better quality with regard to binding data, crystal structures and the nature of the complexes.

In this paper, we use the term **bindability**^15,26^ to describe pockets that bind drug-like ligands with submicromolar affinity and refer to the positive and negative instances in the DaPB dataset as **bindable** and **less-bindable** pockets, in order to distinguish the concepts of druggability and bindability in the NRDLD and DaPB datasets. Although it was stated that borderline druggable proteins are bindable,^27^ the distinction between bindability and druggability is rather obscure and the two terms are sometimes used synonymously.^5,6^ Thus, the DaPB dataset used in this work was built through filtering or selection of data on the basis of bindability. To collect positive complexes, we extracted the properties of the ligands using Pybel, a Python wrapper for the OpenBabel cheminformatics toolkit, ^28^ and then selected the bindable pockets if they bind ligands that satisfy drug-like properties as defined in Sheridan et al. ^15^ with a binding affinity stronger than 1 μM (Table S1). For the negative instances in the DaPB dataset, instead of introducing a set of rules for filtering out the negative (less-druggable) data, we directly applied one of the models trained on the NRDLD training set, TRAPP-SVM (see below), on the remaining unlabeled data and selected the negative data according to its prediction. In summary, the DaPB dataset used here contained 892 positive instances and 1180 negative instances. One fifth of the DaPB dataset was used as the test set and contained 190 positive instances and 224 negative instances.

### Input data generation

Two types of input format were generated during the TRAPP-pocket procedure as explained in more detail in Ref. 29.

#### Preprocessing of the PDB files

Before interpolating the binding pocket on to a grid and computing descriptors, the input PDB file of the protein was preprocessed in two steps. First, redundant small molecules such as solvent, buffer, or some peptide linking groups were removed if they were not in the list of retained co-factors (Table S2). The protein residues and retained co-factors were then protonated at pH 7.0 using Pybel.^28^

#### Multi-channel grid

8 different 3-dimensional grids, each corresponding to a different channel as shown in Table 1, were generated for each pocket. Each channel describes a specific property of the pocket (Figure 1), resembling the idea of a color channel in an RGB image. The size of the grid box can be defined by either the grid edge length, e.g. 24 Å, or the extent around the ligand. For druggability assessment, only a fixed grid edge length was used, due to the limitation of a fixed input dimension for the CNN model. In this 4-dimensional grid representation of the pocket, the first channel describes the shape and position of the cavity, which is computed using the cavity distribution function *G*(**r_i_**, *p*) as described in Ref. 29 with a new pocket selection algorithm summarized in the Supplementary Information. For a cavity grid in which a single grid point is denoted as *r_i_* and the protein as *p*, an atom-occupied position has the grid value *G*(**r_i_**, *p*) = 0 and a pocket position has the grid value 0 *< G*(**r_i_**, *p*) ≤ 1. The remaining 7 channels describe the physicochemical properties of the cavity and were assigned in two steps. First, the atoms in the grid box were assigned to certain physicochemical properties if they satisfied the criteria given in Table 1. The positively and negatively charged atoms were assigned based on the atom and residue information, whereas the atoms with other properties were specified using Pybel ^28^ definitions. Second, Gaussian distribution functions *G^prop^*(**r_i_**, *p*) were spanned over those atoms with certain properties in order to map those atomic features to the grid.

**Table 1:**
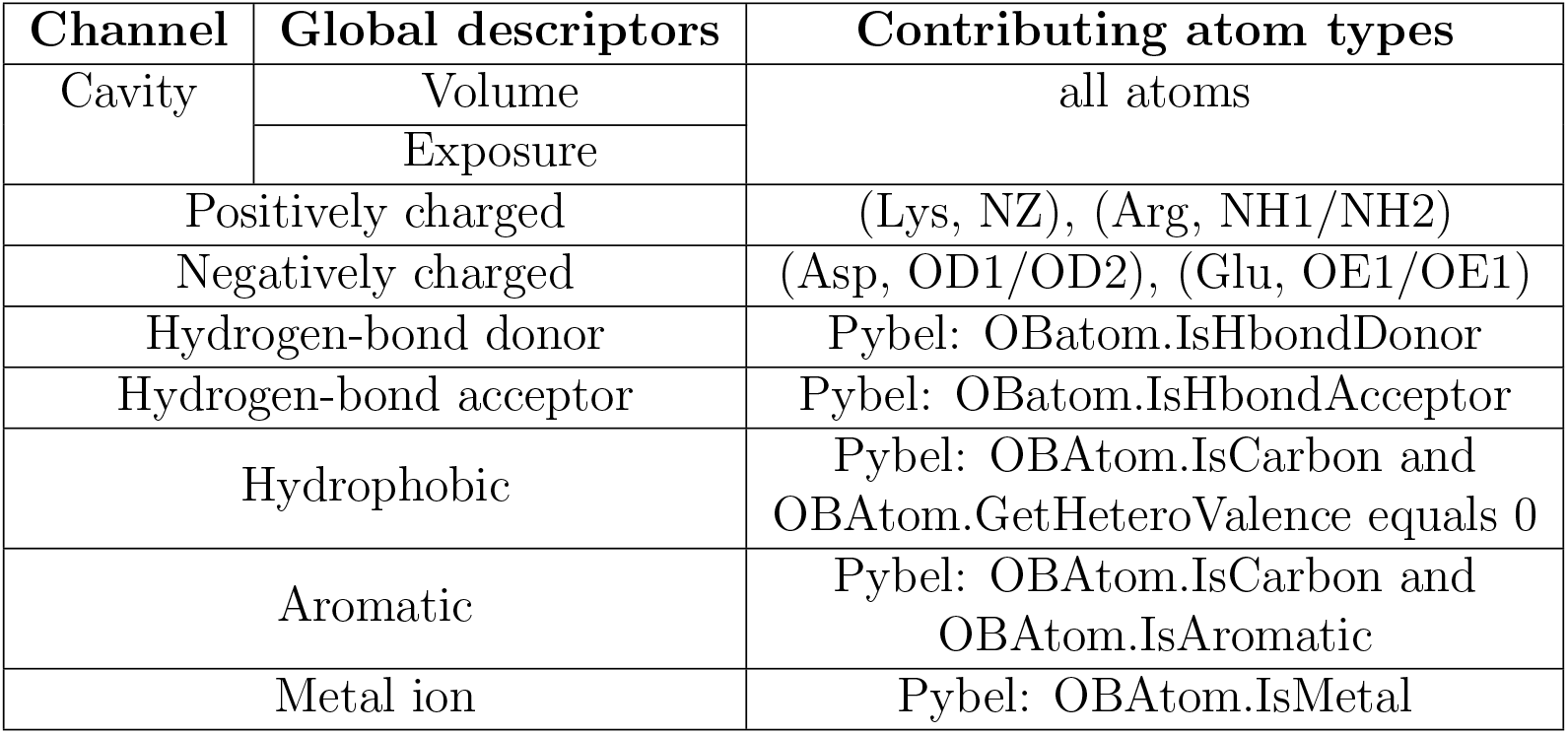
List of 3-dimensional grids and their corresponding global descriptors generated from TRAPP-pocket for the 8 channels.

**Figure 1:**
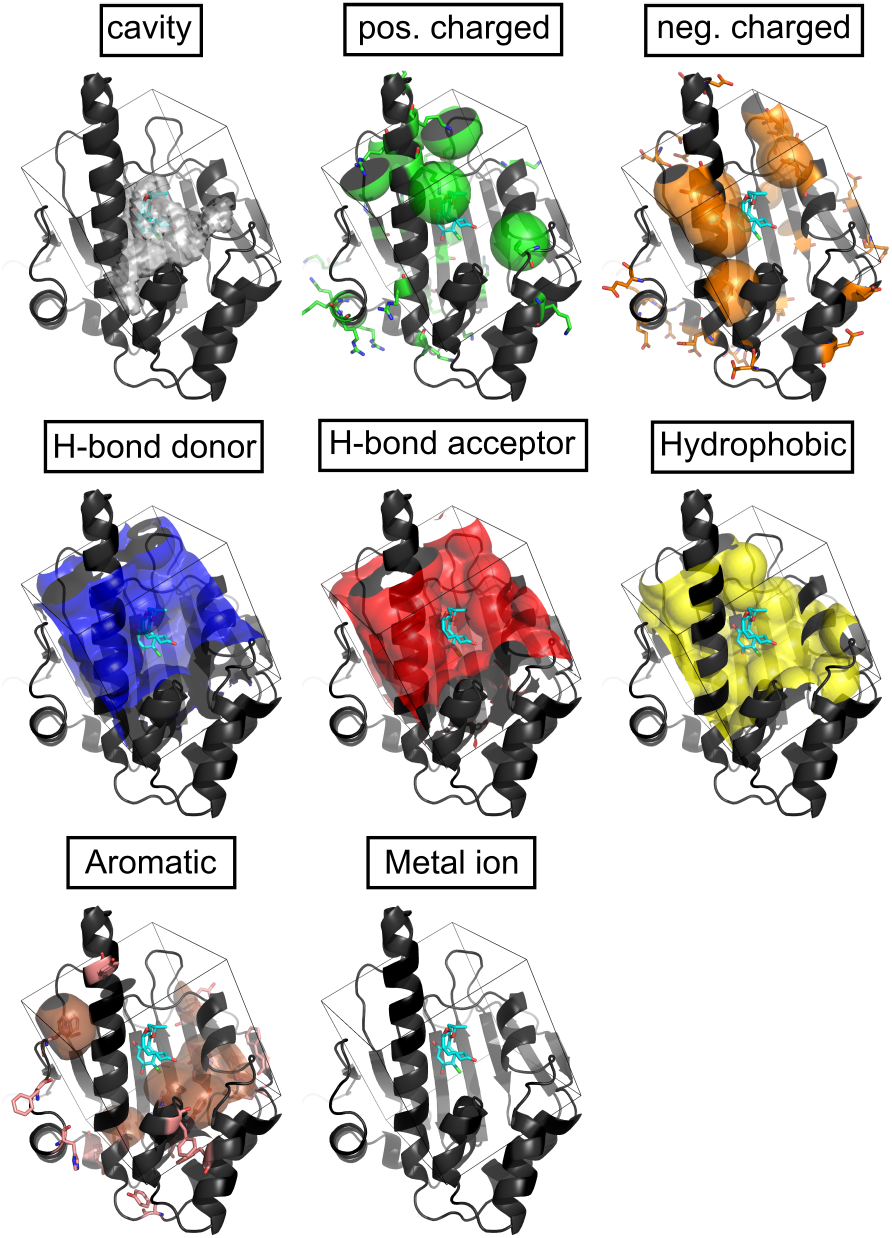
Visualization of the eight channels in grid representation for the structure of radicicol-bound Hsp90 (PDB ID: 1bgq). Radicicol (shown in cyan) is used only for defining the center of the pocket and is removed during grid generation. The grid edge length and grid spacing are 24 Å and 0.75 Å, respectively. The isosurfaces are shown at a level of 0.1. The metal ion channel is empty since there is no metal ion in the binding pocket of Hsp90.

#### Global descriptors

After the grid computations, each channel was compiled into a single value or a few values describing the global pocket properties. In total, ten global descriptors were extracted from the grids in the TRAPP-pocket procedure. The list of the ten global descriptors and their definitions are shown in Table S4. The pocket volume, protein-exposed and solvent-exposed surface area were computed from the cavity grid. The pocket volume is defined as the summation of the grid points that have *G*(**r_i_**, *p*) > 0 and then multiplied by the volume of a single grid element. The protein-exposed and solvent-exposed surface areas were computed by counting the number of transitions from a cavity grid point (*G*(**r_i_**, *p*) *>* 0) to the protein grid point (*G*(**r_i_**, *p*) = 0) or solvent grid point (*G*(**r_i_**, *p*) = −1), respectively, and then multiplying by the square of the grid spacing. The ratio of solvent-exposed surface area to protein-exposed surface area is defined as the pocket exposure, which is an approximate measure of the pocket shape and is used as one of the global descriptors for druggability prediction.

The other physicochemical property descriptors were computed based on the overlap between the cavity grid and the corresponding property grid. If *G*(**r_i_**, *p*) > 0, the property is incremented by *G^prop^*(**r_i_**, *p*). The final value representing each property was then multiplied by the volume of a single grid element in order to make it comparable to the other descriptors that are derived with a different grid setting.

### Statistical Protocol

#### Linear models based on global descriptors

The global pocket descriptors were taken as the inputs for the druggability assessment models using logistic regression and linear SVM, as implemented in scikit-learn.^30^ The models, whose number of parameters is the number of input features plus one for the bias term, were trained on the small NRDLD training set with L1 or L2 weight regularization. The pipeline, which consisted of input generation, data standardization, and model prediction, was constructed and optimized as a whole. The hyperparameters, e.g. grid size and grid spacing for grid generation, as well as the parameter C, which controls the regularization strengths, were optimized by a 5-fold cross-validated grid search (Table S5). The splitting of the validation set was performed so as to ensure that each fold had similar numbers of the instances with the two labels.

#### Convolutional neural network based on a multi-channel grid

To capture the relationships between the druggability and the spatial/physicochemical properties of the binding pockets, the multi-channel grid, which serves as a detailed description of the binding site, was used as the input for druggability prediction with a convolutional neural network. The network architecture used for druggability prediction was inspired by Ref. 31, which had 3 3 × 3 × 3 convolutional layers with rectified linear activation units alternating with max pooling layers followed by a fully connected layer with two outputs and a softmax layer to obtain a probability distribution over two classes.

The DaPB dataset was used for training and hyperparameter searching of the neural network. To combat overfitting, data augmentation through 90° rotation and random translation of 2 voxels in *x, y, z* coordinates for each training instance was performed. The network architecture and other hyperparameters, such as the learning rate and the weight decay for regularization, were optimized using a 3-fold cross-validated grid search. In total, 60 different network architectures with varying numbers of convolutional layers, numbers of convolutional filters per layer, and numbers of neurons in the fully-connected layer were compared based on their cross-validated F1 score. Finally, the optimized network structure was trained using the Adam optimizer with default parameters for momentum scheduling (*β*_1_ = 0.99, *β*_2_ = 0.999), a learning rate of 0.001 and a weight decay of 0.0001.

#### Model evaluation

The quality of the models was assessed using accuracy (Eq. 1), sensitivity, specificity (Eq. 2), and Matthew’s correlation coefficient (MCC, Eq. 3). These metrics can be derived based on the four elements in the confusion matrix during binary classification, that is, the numbers of true positives (TP), true negatives (TN), false positives (FP), and false negatives (FN). The sensitivity and specificity indicate the ability to identify druggable and less-druggable pockets, respectively. The MCC, which takes into account all four values, is a balanced metric for an uneven class distribution and an ideal measure for assessing the quality of a binary classifier. Since the aim is to achieve a good balance between precision and recall, the harmonic mean of precision and recall, also known as the F_1_ score or F-measure, is used in the cross validated grid search for the optimal hyperparameters (Eq. 4).

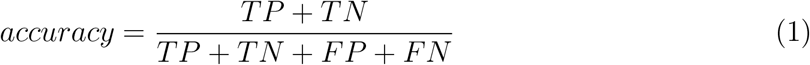

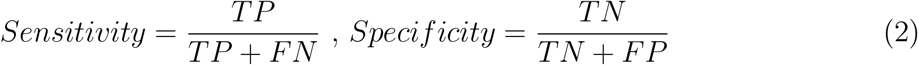

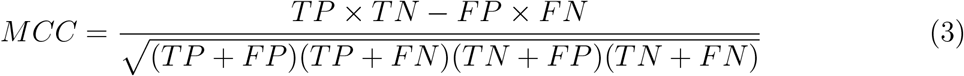

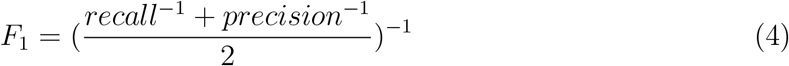

### Interpretation and visualization of the druggability prediction models

To interpret the results of the linear models, the property values and the contribution of each property to the decision were visualized as a heatmap. By displaying the standardized input values, as well as color-coding each global descriptor by the product of the input value and the corresponding weight, the dominating features were identified.

On the other hand, the decision of a CNN can be explained by layer-wise relevance propagation (LRP),^32^ which is a technique to calculate the contribution of each grid point in the input space to the prediction made by the network. The relevance, which is initialized as the network output before the softmax operation, is propagated back through the network until it reaches the input. Through propagating relevance proportional to the pre-activation of a specific node at the previous layer, the conservation of relevance at each layer can be realized. The LRP result is also a grid with the same dimension as the input grid but each grid point stores the relevance or contribution of that position to the output, thus the regions in the binding pocket with a strong contribution to the model prediction can be visualized as a 3D mesh. Since the output layer of TRAPP-CNN contains two units representing druggable and less-druggable pockets, LRP can be performed from both units in order to find the critical regions that make the pocket druggable or less-druggable. However, as the relevance of each grid point is a decomposition of the output, the magnitude of the relevance depends on both the importance of that position to the prediction and the magnitude of the output, which represents how certain the model is of the prediction. In order to visualize the LRP results as 3-dimensional isocontours, the relevance at each grid point was first normalized by the maximum relevance value in the grid and an empirical threshold of 0.1 was used to show only the regions with a significant contribution. The sub-regions of the pocket showing a strong contribution to the model decision as druggable or less-druggable were colored red or blue, respectively.

#### Model testing using a trajectory from an L-RIP simulation

To assess the ability of our models to monitor druggability in an MD simulation, we used an enhanced sampling MD method called L-RIP^33^ (Rotamerically Induced Perturbation ^34^ with Langevin dynamics) to sample different conformations of the active site of p38 mitogen-activated protein (MAP) kinase and the cryptic pocket of TEM-1 *β*-lactamase. The p38 MAP kinase is an intracellular signaling protein involved in cytokine synthesis and is an important target for the treatment of osteoarthritis and inflammation. *β*-lactamase is a hydrolase that breaks the *β*-lactam ring, leading to resistance to *β*-lactam antibiotics among bacteria. To study how transient pocket formation in these two example proteins impacts druggability, L-RIP perturbation was applied with 300 perturbation pulses and 300 implicit solvent MD timesteps of 0.002 ps after each perturbation. ^33^ The last frame of each perturbation step was stored for further analysis. The remaining parameters for applying L-RIP followed the default settings on the TRAPP webserver.^33,35^ Before druggability analysis, all snapshots in the trajectory were superimposed by fitting all binding site heavy atoms that were within 4 Å of the centre of any ligand atom.

## Results

### Three global descriptors - pocket volume, hydrophobicity and the number of H-bond donors - are the most important for predicting druggability using linear models

Before training the models, we first inspected the data sets by plotting the global descriptors in a bar chart and a scatter-plot matrix (Figure 2A and B). We observed similar trends in both the NRDLD and the DaPB datasets, indicating that both sets convey consistent information regarding the relationship between druggability and the global descriptors. Comparing the mean of each global property within the DaPB set (Fig. 2A), we observe that the values for the pocket volume, number of H-bond acceptors, hydrophobicity and aromaticity of the druggable targets are higher than those of the less-druggable targets. In contrast, the values for the pocket exposure, the positively charged property, and the number of H-bond donors of the druggable targets are slightly lower than those of the less-druggable targets. The large standard deviation with respect to the mean for each global property for both druggable and less-druggable pockets demonstrates the variability of different protein binding pockets.

**Figure 2:**
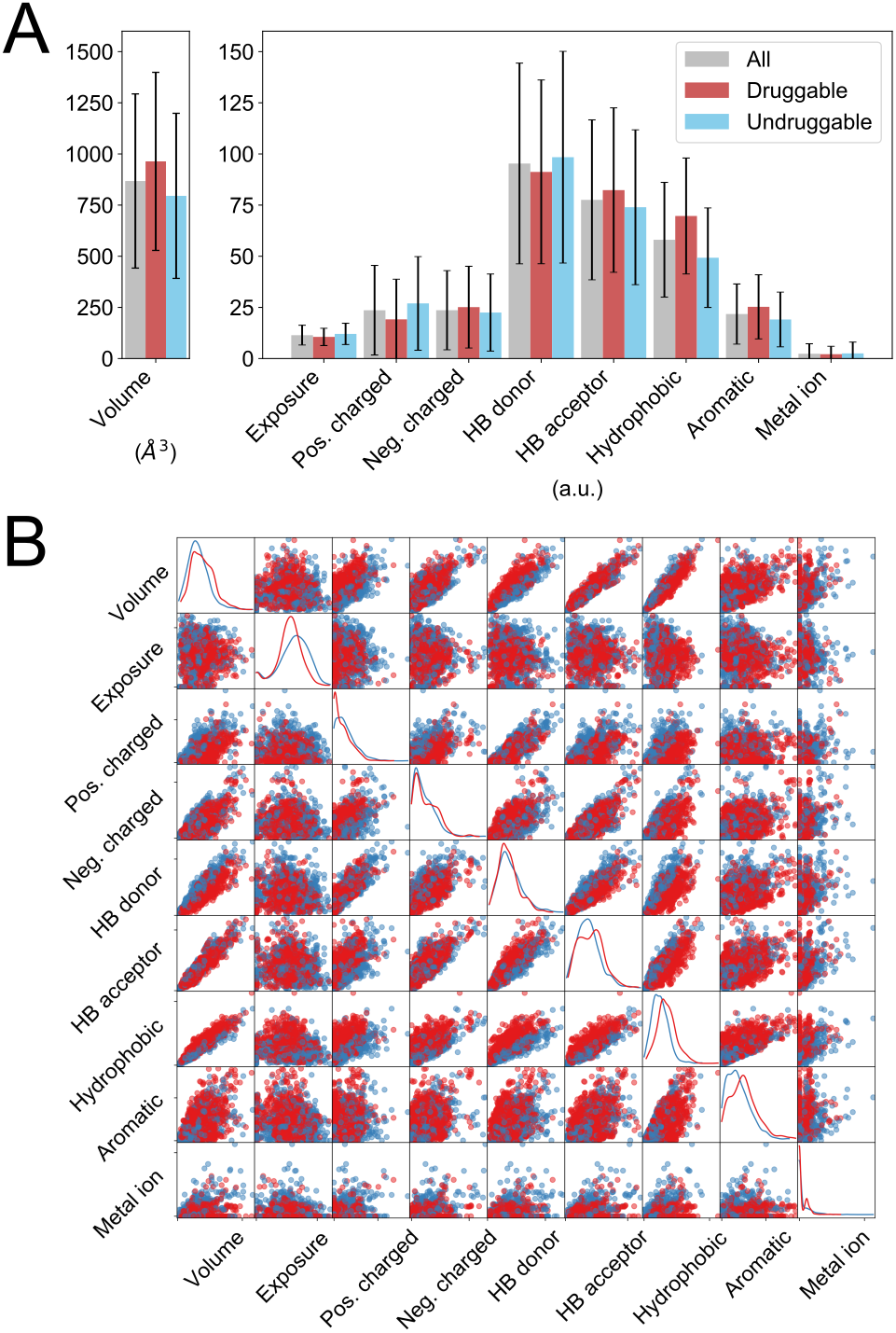
Visualization of the DaPB dataset built on the PDBbind refined set 2017. (A) Mean and standard deviation of each global property computed using all, bindable and less-bindable protein structures in the DaPB dataset. (B) Scatter-plot matrix of the DaPB dataset. The red and blue dots represent bindable and less-bindable binding pockets, respectively. The corresponding plot for the NRDLD set is given in Figure S1

On the other hand, the scatter-plot matrix allowed us to detect correlations between pairs of global features (Figure 2B). First, we observed a linear correlation between the pocket volume and some of the physicochemical properties, such as the numbers of H-bond donors and acceptors and the hydrophobicity. This correlation reflects the fact that the computation of a physicochemical property is basically a summation of certain atom types located within the pocket and thus the value is proportional to the pocket volume. We therefore included the normalization of each physicochemical property to the pocket volume as one of the hyperparameters (parameter values: “yes”/“no”) to be tuned during optimization. It was noticeable that the scatter plots for hydrophobicity/positively charged or hydrophobicity/number of H-bond donors show some linear separability between bindable and less bindable pockets, indicating the importance of these global features. In contrast, the most severe overlaps between the positive and negative data for bindability occur for the scatter-plots for the pocket exposure, negatively charged, and aromaticity features.

To construct linear models using LR or a linear SVM, the global features extracted from the TRAPP procedure were used as input for classification. Both methods are simple, fast, and ideal for binary classification, and less prone to over-fitting for small datasets than more complex machine learning methods. In addition, the linear dependency of the models on the global features provides good interpretability. The hyperparameters of the final model that achieved the top cross-validated score on the NRDLD training set were a grid size of 24 Å, a grid spacing of 0.75 Å, and *C* = 1 with the physicochemical properties normalized by the pocket volume.

The resulting coefficients of the TRAPP-LR with L1 and L2 regularization and TRAPP-SVM models are similar (Figure 3). We first consider the interpretation of the coefficients obtained in the L2 regularized TRAPP-LR model. The output of LR is the log-odds ratio. Thus, positive coefficients, e.g. for pocket volume and hydrophobicity, imply that the odds of the pocket being druggable increase as the corresponding input feature increases, whereas negative coefficients indicate that the odds of the pocket being druggable decrease as the corresponding feature increases. Thus, by increasing the pocket volume and hydrophobic groups in the binding pocket while keeping the other features unchanged, the binding pocket would be more likely to become a drug target. In contrast, larger values for the number of H-bond donor/acceptor groups, positive charged groups, and the metal ion property lead to a lower probability that a pocket is druggable.

**Figure 3:**
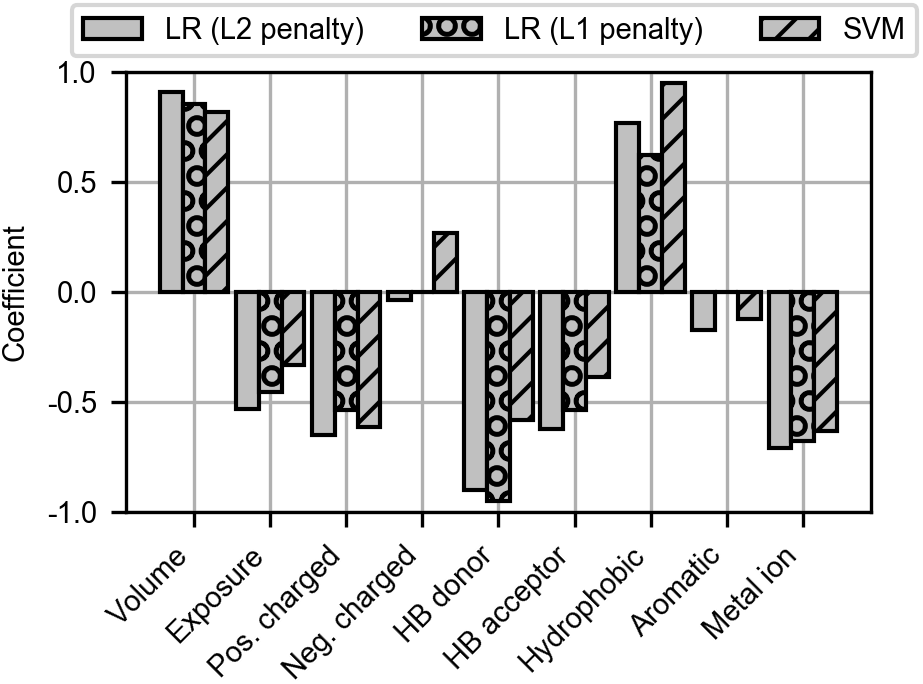
Coefficients of the descriptors in the TRAPP-LR models with L2 (grey boxes) and L1 (circle hatch) regularization and the TRAPP-SVM model (line hatch) trained on the NRDLD training set. All models have input descriptors on grids with a grid edge length of 24Å, a grid spacing of 0.75Å, a hyperparameter C of 1, and normalization of physicochemical properties to the pocket volume.

For all three models, the pocket volume and hydrophobicity are the main properties that are positively correlated with the druggability, while the other global properties mostly have a negative correlation. The importance of each feature can be derived from the absolute size of the coefficient relative to the others. We observe that some features, including pocket volume, positively charged, number of H-bond donors and hydrophobicity, consistently have larger coefficients in all models. Notably, the metal ion property contributes more than several other physicochemical properties, such as aromaticity and the number of H-bond acceptors, which are commonly included in druggability predictions. Note that most of the metal ions in the NRDLD dataset have a formal charge of +2e (Table S3). Geometric descriptors have been widely used for druggability prediction, for example, the pocket depth or buriedness has been shown to have a positive contribution.^11,14,15,36^ However, we find that the coefficient of pocket exposure is relatively small. This difference may arise because some of the previous models were built using only a few simple input features, thus overestimating the importance of pocket shape. ^15,36^ The use of a different druggability training set may also affect the coefficients of the final model.^11^ Another possible reason is that the ratio between solvent-exposed and protein-exposed surface area is not an ideal shape descriptor. In PockDrug,^14^ the longest distance in the pocket or the radius of the smallest enclosing sphere were used instead.

The coefficients of the TRAPP-LR models with L1 or L2 regularization were overall similar. L1 regularization penalizes the absolute value of the weights instead of the squared value penalized in L2 regularization. As the L1 penalty induces sparsity, the weights corresponding to the negatively charged descriptor and aromaticity vanished, indicating that these two properties are less important for druggability prediction. This is as expected because of the severe overlap of the druggable and less-druggable classes in these two feature dimensions (Figure. 2B).

### Large hydrophobic patches in the binding pockets contribute strongly to the druggability prediction of the TRAPP-CNN model

The DaPB training set was used for training the TRAPP-CNN. To be consistent with the procedures for TRAPP-LR and TRAPP-SVM, the same configuration for grid generation, that is, a grid size of 24 Å and a grid spacing of 0.75 Å, was used. Based on the network architecture of Ref.,^31^ we further tuned our network architecture by grid searching over networks with varying depth and width (Tables S6 and S7). In general, increasing the number of convolutional layers slightly degraded the performance. Modifying the number of convolutional filters in each layer did not show an obvious effect but adding an additional fully-connected layer did improve the performance. The final network architecture is shown in Figure 4.

**Figure 4:**
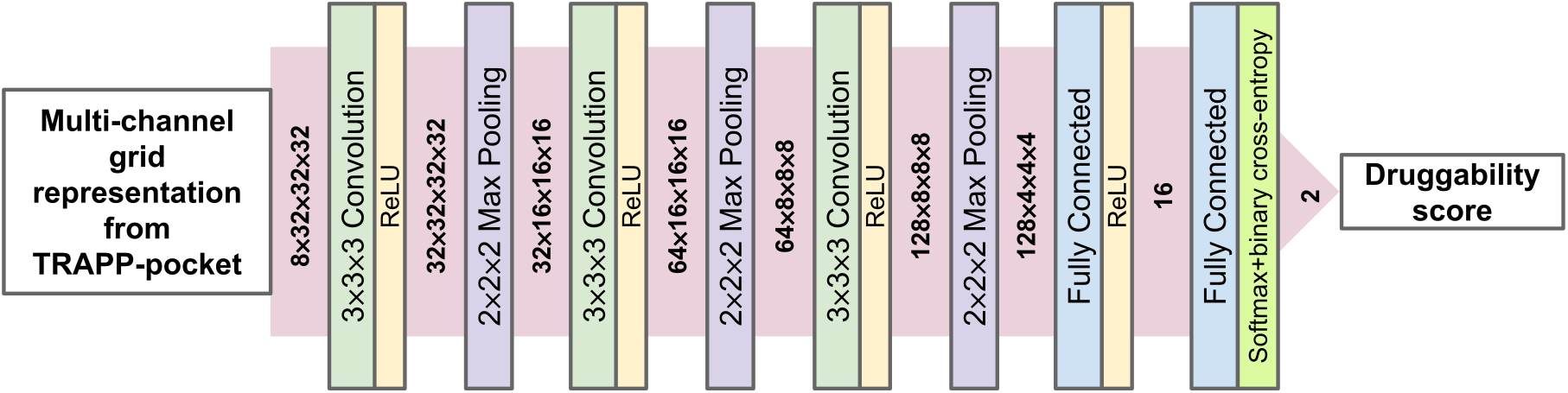
Final network architecture of TRAPP-CNN.

To provide better interpretability, the decisions of TRAPP-LR and TRAPP-CNN were visualized with a heatmap and by pixel-wise decomposition through LRP, respectively. Two positive and two negative examples in the DaPB test set that are correctly predicted by both TRAPP-LR and TRAPP-CNN are shown in Figure 5. On the heatmap, the standardized global property values obtained after preprocessing are displayed as numbers, providing information on how the global properties of a given pocket deviate from an average pocket. The druggability score of TRAPP-LR is the inner product of the standardized global properties and the weights plus a bias term. The contribution of each property can therefore be derived from the product of the property value and its corresponding weight. The contributions from different properties are displayed by the color of each property. The stronger the red color, the more positive the contribution of a certain property to the prediction. In contrast, a stronger blue color indicates a more negative contribution. Thus, a generally red matrix indicates high druggability, whereas an overall blue matrix indicates low druggability. Note that the properties are ordered according to the magnitude of the weight, therefore the more important features are shown in the upper left corner and the less significant features are in the lower right corner.

**Figure 5:**
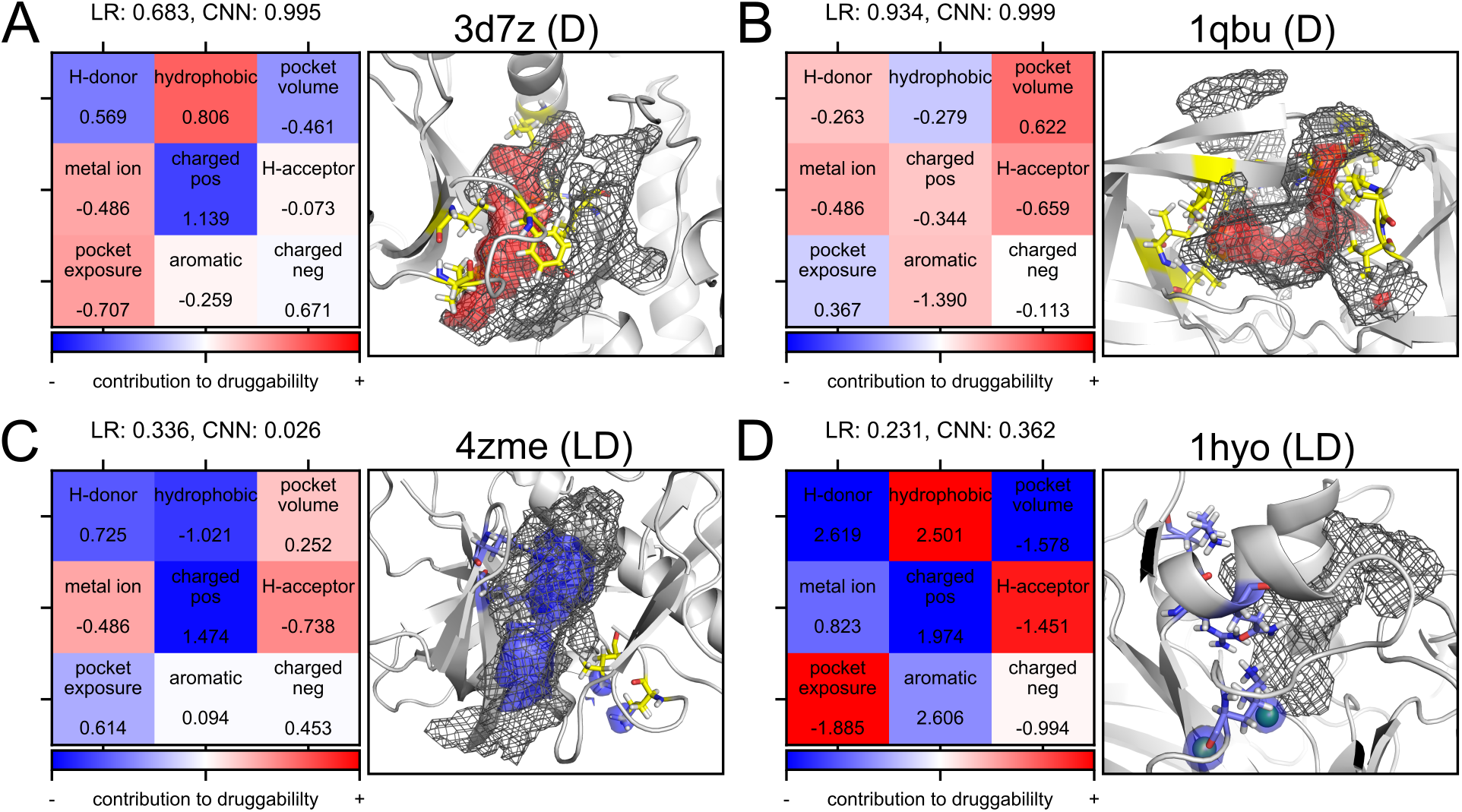
Heatmaps from the TRAPP-LR model and pixel-wise decompositions from the TRAPP-CNN model for two positive and two negative examples in the DaPB test set. The PDB ID, druggability label, and its predicted druggability scores are shown above the panels for each example. The druggability labels are shown as **D** for druggable and **LD** for less druggable. In the heatmap, the scaled input values are shown in numbers below each property name, while the contribution from each property is shown by the color (red-druggable, blue - less druggable). The cavity extracted by TRAPP-pocket is shown as a black mesh. Evidence for druggable and less-druggable pockets is shown by red and blue isosurfaces, respectively. The residues shown in yellow are hydrophobic and the residues in blue are positively charged. Metal ions are shown as green spheres. (A) p38 MAP kinase (3d7z). (B) HIV protease (1qbu). (C) Myosin-II Heavy Chain Kinase A (4zme). (D) Fumarylacetoacetate hydrolase (1hyo).

The LRP results of TRAPP-CNN can be visualized as meshes on the structure of a binding pocket in order to identify critical sub-regions of the pocket for its bindability. The positive and negative evidence are shown by red and blue regions, respectively. In general, we observe that most of the evidence is located close to or within the pocket mesh, demonstrating that the cavity grid generated from TRAPP-pocket is able to guide the model to the region of interest. On the other hand, for pockets predicted to be in the positive class (druggable), mainly positive evidence but not negative evidence is obtained and vice versa. This phenomenon is especially prominent for the pockets with predictions of very high confidence, that is, druggability scores close to 0 or 1. As pixel-wise decomposition distributes the model raw output onto the input grid points, a confident prediction is associated with a large difference between the output of the two classes. Therefore, the evidence corresponding to the class opposite to the prediction should be negligible.

Two positive and two negative examples in the DaPB test set are shown (Figure 5). Comparing the two positive examples (Figure 5A and B), the heatmap of 1qbu (closed conformation of the active site of human HIV protease) contains relatively more red fields than that of 3d7z (ATP binding site of human p38 MAP kinase), which corresponds to the difference in druggability scores predicted by TRAPP-LR. The large hydrophobicity of the pocket in structure 3d7z is the main contributing factor, while the large pocket volume and the relatively H-bond donor/acceptor properties of the poicket in the structure 1qbu make it a druggable target. The LRP results of the 3d7z and 1qbu structures demonstrate that the positive evidence is situated in the main pocket and encompasses most of the pocket mesh, indicating that the cavity volume has the largest contribution to the positive (druggable) prediction. A closer examination of the binding site residues reveals that the positive evidence tends to be surrounded by hydrophobic residues. Thus, hydrophobicity is a critical feature for enabling TRAPP-CNN to discriminate druggable pockets from less druggable ones.

The negative examples shown are the *α*-kinase domain of myosin-II heavy chain kinase A and fumarylacetoacetate hydrolase (FAH). The *α*-kinases (PDB ID:4zme, Figure 5C) are serine/threonine protein kinases that exhibit no sequence similarity with conventional eukaryotic protein kinases.^37^ The active site of *α*-kinase is, compared to that of p38 MAP kinase, more solvent-exposed and less hydrophobic according to the heatmaps. Two regions with negative evidence are observed. One of them is surrounded by positively charged residues, which corresponds to the large negative coefficients for the positively-charged descriptor in the TRAPP-LR and TRAPP-SVM models. The other region with negative evidence is within the pocket mesh and overlaps with the position of the bound adenosine in a rather shallow sub-region of the pocket. Note that some additional negative evidence is also present near the side chains of two leucine residues that are not pointing towards the binding pocket. Hence, the noncontributing hydrophobic residues near the binding pocket are also recognized by the model as part of the reason for low druggability.

The binding site of FAH (PDB ID: 1hyo, Figure 5D) is an example of a protein pocket that contains metal ions. The heatmap for 1hyo indicates that the small pocket volume and relatively higher H-bond donor/ positively charged features are the key reasons for its low druggability. Interestingly, the negative evidence near the binding site of FAH is concentrated on the metal ions, a calcium ion and a magnesium ion, which can also be understood from the negative coefficient for the metal ion property in the TRAPP-LR model. If we repeat LRP on the same reference structure with no metal ion, the prediction remains as negative (less-druggable) but the negative evidence transfers to the center of the narrow cavity, surrounded by positively charged residues, as well as a shallow region of the pocket. The druggability score of FAH from TRAPP-LR increases from 0.231 to 0.343 upon removal of the metal ion, showing the direct negative contribution of metal ions to the druggability. In contrast, the druggability score from TRAPP-CNN decreases slightly from 0.362 to 0.316, indicating that, in reality, the impact of metal ions on the prediction is not as severe as indicated in the visualization. To sum up, the visualization through LRP might lead to over exaggeration of the negative signal from metal ions, and therefore, druggability prediction for protein pockets containing metal ions should be interpreted with caution and repeated using the metal ion-free structures.

### The TRAPP-based druggability prediction models show equivalent or slightly improved performance on a public test set

The performance of the tuned models was first evaluated using the DaPB test set, which is the 1/5th of the DaPB dataset that was not used for training, see Table 2. As the negative data were labeled by the TRAPP-SVM model, its specificity is 1 and both accuracy and MCC are high. The only informative metric for the TRAPP-SVM model is the sensitivity (true positive rate), which implies a difference between bindable pockets, defined by filtering drug-like properties with high affinity, and druggable pockets described in the NRDLD dataset. Only 72.5% of the positive instances in the DaPB dataset were recognized as druggable by the TRAPP-SVM model. Having a similar decision boundary according to the coefficients of the model (Figure 3), the TRAPP-LR model shows comparable performance to the TRAPP-SVM model with similar sensitivity on the DaPB test set. It can be that the misclassified positive instances are less-druggable but bindable pockets based on the definition of druggability in the NRDLD dataset. Interestingly, the TRAPP-CNN model, which is the only model trained on the DaPB dataset, performed similarly to the other two linear models with higher specificity than sensitivity. This indicates that the two classes might still be overlapping in the convolved feature space. As stated above, the positive instances might contain less-druggable but bindable instances. Since the positive instances are selected using observation-based rules for drug-like properties which are strongly dependent on the ligand properties, there might still be bindable pockets in the rest of the PDBbind refined set. The negative data that are chosen from the rest of the PDBbind refined set based on the prediction of the TRAPP-SVM model might thus also contain less-druggable but bindable pockets. These data points in the convolved space, which is optimal for separating druggable from less-druggable pockets, are likely to be close and in the vicinity of the decision boundary. As the negative data are slightly more numerous than the positive data, the model would be optimized more towards correct classification of the less-druggable class, leading to the higher specificity than sensitivity (Figure S2A, S2B).

**Table 2:** Predictive performance of the TRAPP-LR/SVM/CNN models for the DaPB and NRDLD test sets and comparison to other druggability prediction tools for the NRDLD test set. The value for CavityDrugScore is from the original publication^16^ while the values for Volsite, DrugPred and PockDrug are from Table 3 in Ref. 14. The values for the TRAPP-SVM model on DaPB test set are shown in parentheses since the TRAPP-SVM model was used for defining the negative data in the DaPB set.

| Dataset | DaPB test set |  |  | NRDLD test set |  |  |  |  |  |  |
| --- | --- | --- | --- | --- | --- | --- | --- | --- | --- | --- |
| Method | TRAPP-SVM | TRAPP-LR | TRAPP-CNN | Cavity-DrugScore | Volsite | DrugPred | PockDrug | TRAPP-SVM | TRAPP-LR | TRAPP-CNN |
| Test accuracy | (0.882) | 0.847 | 0.838 | 0.82 | 0.89 | 0.89 | 0.865 | 0.892 | 0.892 | 0.946 |
| Test sensitivity | (0.725) | 0.713 | 0.742 | - | - | - | 0.957 | 0.913 | 0.913 | 0.913 |
| Test specificity | (1.000) | 0.949 | 0.920 | - | - | - | 0.714 | 0.857 | 0.857 | 1.000 |
| Test MCC | (0.775) | 0.694 | 0.678 | - | 0.77 | 0.77 | 0.712 | 0.770 | 0.770 | 0.894 |

To compare with other existing druggability prediction tools, the performance of all three models was further evaluated on the NRDLD test set as shown in Table 2. The TRAPP-LR and TRAPP-SVM models perform equally to the other methods in terms of accuracy and MCC value. The TRAPP-CNN model, however, displays superior performance with only 2 misclassified positive data points. Note that the number of data points in the test set is low (37 instances), and thus the difference in performance arises from a difference in the misclassification of just two data points.

To interpret the predictions and figure out the reasons for the wrong predictions, we examined the misclassified examples from the TRAPP-LR model (Figure 6). Note that the druggability score for the TRAPP-LR and TRAPP-CNN models can be interpreted as the posterior probability (Figure S2C, S2D). The decision is given by thresholding at 0.5. As the NRDLD data set contains more positive examples than negative examples, the prior probability of the positive class, which is part of the posterior probability, is overestimated and could be corrected. However, since the goal is not to obtain a binary prediction but to compare the scores between different conformations, we can keep the score unmodified but instead use a higher threshold for selecting promising conformations. In these misclassified examples, if the threshold is increased to 0.61, the false positive examples would be eliminated.

**Figure 6:**
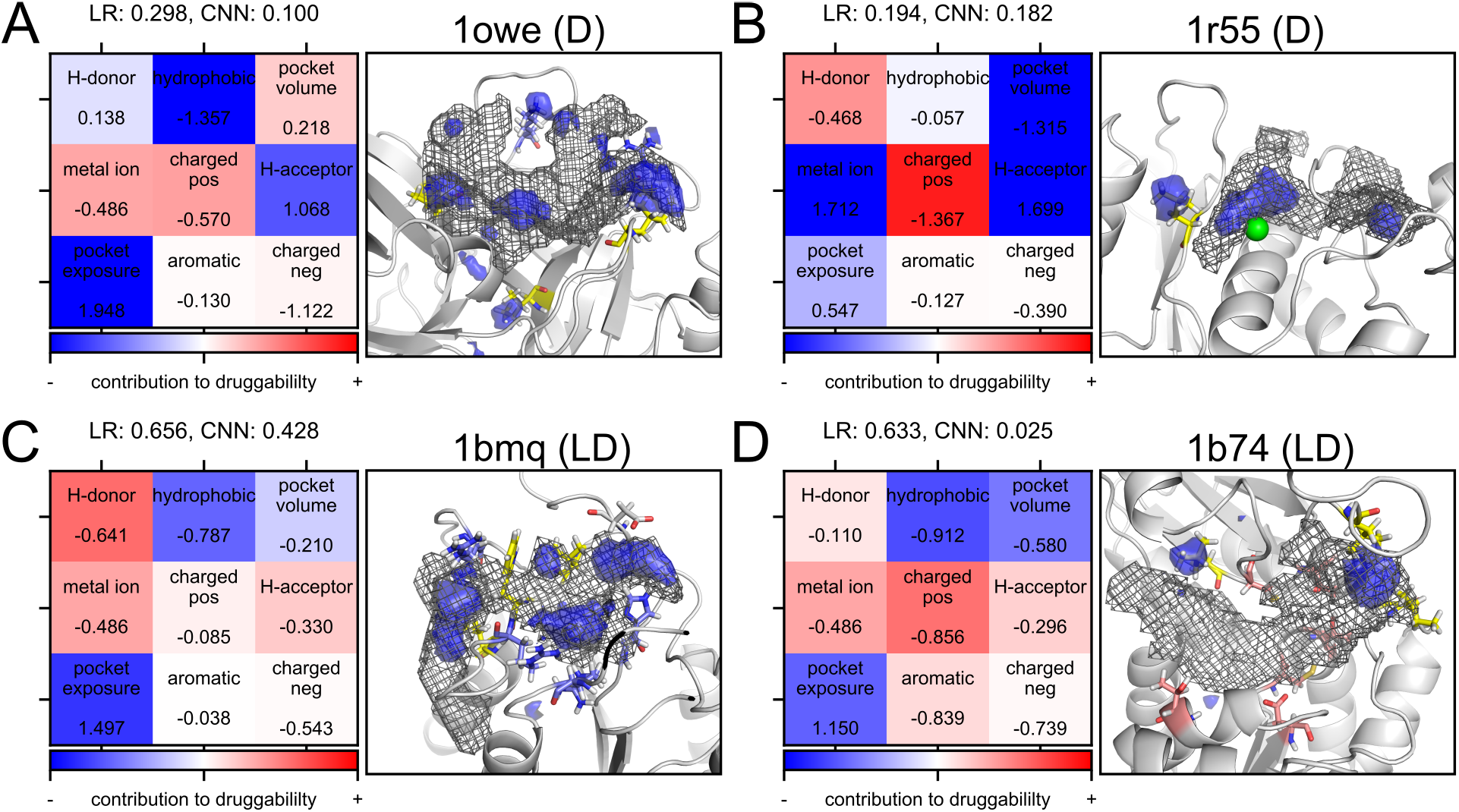
False negatives and false positives in the NRDLD test set from the predictions of the TRAPP-LR model. The PDB ID, druggability label, and its predicted druggability score from the TRAPP-LR model are shown above the panels for each case. **D** indicates druggable and **LD** indicates less druggable according to the NRDLD data set. The cavity extracted by TRAPP-pocket is shown as a black mesh. Evidence for druggable and less-druggable pockets is shown by red and blue isosurfaces, respectively. The residues shown in yellow, blue, and pink are hydrophobic, positively charged, and H-bond forming residues. Metal ions are shown as green spheres. False negatives: (A) urokinase (1owe). (B) ADAM 33 (a disintegrin and metalloprotease, 1r55). False positives: (C) interleukin-1 beta converting enzyme (1bmq). (D) glutamate racemase (1b74).

In terms of the heatmaps, the misclassified examples, especially 1bmq and 1b74, have less strong color than the examples in Figure 5, indicating that none of these properties makes significant contributions to the predictions of the model. This is another sign of the lack of confidence in the predictions. For the urokinase (PDB ID: 1owe), the low druggability comes from the low hydrophobicity, additional H-bond acceptor groups and an overly solvent-exposed environment. The binding pocket of ADAM 33 (a disintegrin and metalloprotease, PDB ID: 1r55) is also predicted as less-druggable due to the smaller pocket size and extra H-bond acceptors. For the interleukin-1 beta converting enzyme (ICE, PDB ID: 1bmq), the lower H-bond donor and acceptor properties of the pocket lead to the prediction of slightly druggable. The cavity of glutamate racemase (PDB ID: 1b74) has a relatively low hydrophobicity and pocket volume, common characteristics of less-druggable pockets. However, the low amount of positively charged groups in the pocket seems to increase the probability to be druggable and reverts the prediction.

We further applied the TRAPP-CNN model to these four examples that were misclassified by the TRAPP-LR model and visualized the evidence by pixel-wise decomposition with LRP (Figure 6). For the complete NRDLD test set, 1owe and 1r55 are the only two misclassified cases when using the TRAPP-CNN model, whereas 1bmq and 1b74 are correctly predicted as less-druggable owing to the higher specificity of the TRAPP-CNN model. According to the structures and the cavity meshes, all four pockets are shallow and solvent-exposed. In general, the evidence for less-druggable pockets highlights the regions that are at the surface or outskirts of each cavity, close to some hydrophobic residues. In comparison to many druggable pockets characterized by a well defined hydrophobic environment, these hydrophobic residues tend to be situated on flexible loop structures and are not clustered together in a confined region; thus they are unable to form a hydrophobic patch for ligand binding that would favour a druggable pocket. In 1owe, the shallow regions, as well as some positively charged residues, are identified as evidence for low druggability. The cavity in 1r55 is situated on the protein surface and the bound ligand is barely buried. As the original pixel-wise decomposition result for 1r55 only emphasizes the metal ion, as discussed previously, the procedure was repeated using the metal-ion free structure as input. The less-druggable evidence also points out the shallow region where the ligand binds, as well as two other sub-regions which may be noise for the druggability prediction. Similar to the previous two cases, the shallow regions or dynamic hydrophobic residues close to the exterior of the pocket are highlighted with less-druggable evidence in 1bmq and 1b74. The cavity centers of 1bmq and 1b74 are surrounded by positively charged and hydrogen bonding groups, respectively, which are not characteristics of druggable pockets. Even though 1b74 is correctly predicted as less druggable by the TRAPP-CNN model and has the lowest druggability score among the four examples, the impact of the hydrogen bonding groups is unexplained by LRP.

### More druggable conformations of transient pockets are identified using the TRAPP procedure

To demonstrate how conformational changes impact druggability, we applied our druggability prediction models to two systems containing cryptic binding sites. ^38–40^ It has been reported that appropriately sized pockets of the biologically relevant drug targets are necessary for the strong binding of drug-sized ligands.^41,42^ For the target proteins that are considered tractable but less-druggable, it is of interest to identify their cryptic sites and to exploit those sites to boost the druggability of the drug targets.^41^

The first example considered is the p38 MAP kinase. It has a deep ATP-binding pocket that contacts a flexible *β*-sheet and two flexible loops. The shifting of the F169 side chain from its buried position (DFG-in conformation) to a position that sterically interferes with ATP binding (DFG-out conformation), results in the opening of a hydrophobic sub-pocket adjacent to the ATP binding site which is exploited by diaryl urea inhibitors. ^43^ Inhibitors bound to the active site of the two conformations were discovered, indicating that both are potentially druggable. Nonetheless, comparing the crystal structures, an extra hydrophobic sub-pocket close to the main pocket tends to be more accessible in the DFG-out conformation, leading to an increased potential for specific ligand binding and thus higher druggability scores than that of the DFG-in conformation (Figure 7A and B).

**Figure 7:**
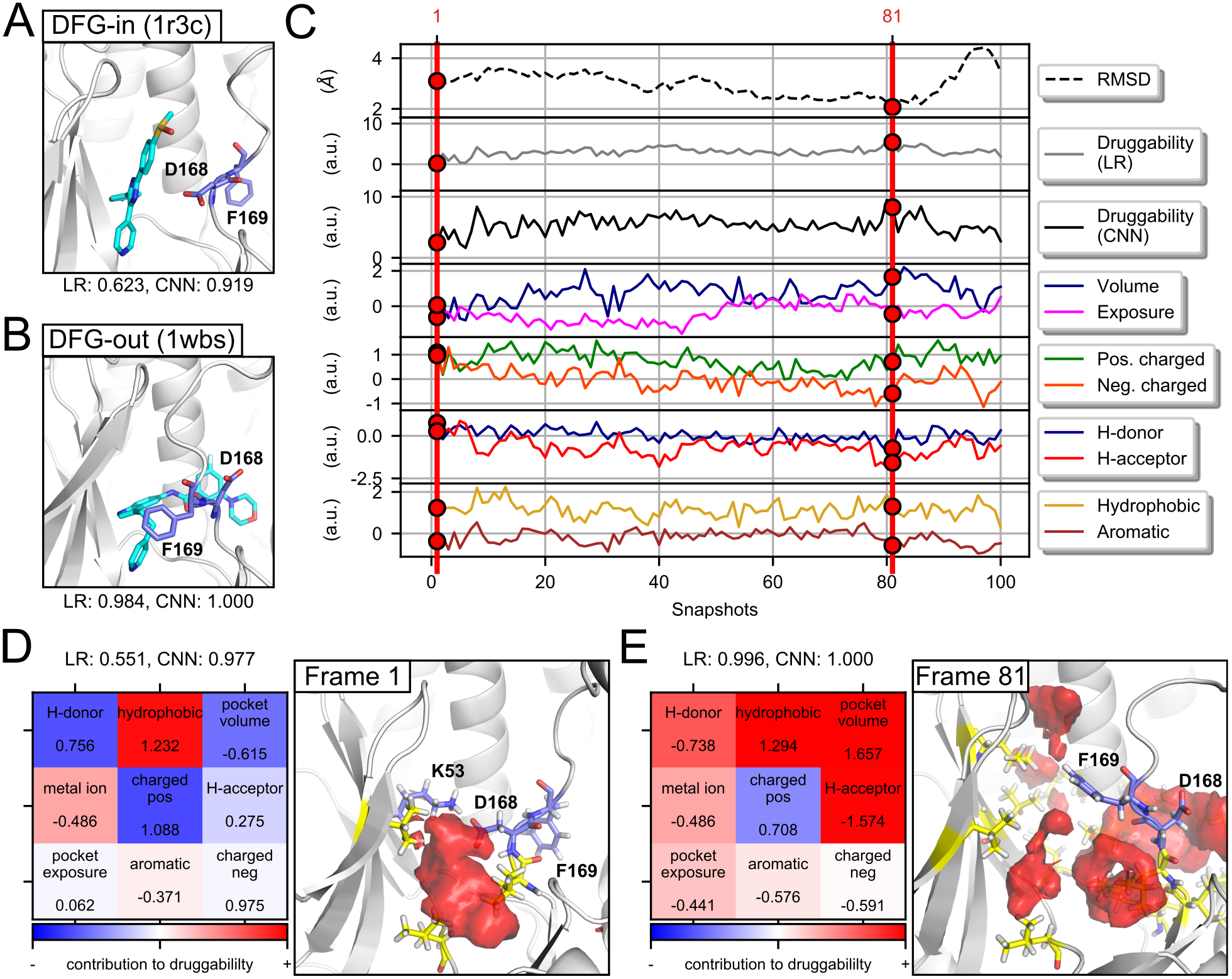
Use of the TRAPP-based models to monitor variations in druggability along an MD simulation of p38 MAP kinase. (A)(B) Crystal structures of p38 MAP kinase in DFG-in and DFG-out conformations. D168 and F169 are shown in blue, while the bound ligands are shown in cyan. (C) Traces of druggability and other physicochemical properties along the L-RIP trajectory starting from the DFG-in conformation (PDB ID: 1r3c, shown in panel (A)). The RMS distance was calculated between each frame and the DFG-out conformation based on the active site residues of p38 MAP kinase (PDB ID: 1wbs, shown in panel (A)). The first frame and the frame 81 that has the lowest RMSD to the DFG-out conformation and the highest predicted druggability, are highlighted. The metal ion property is not shown because there is no metal ion in the binding pocket of p38 MAP kinase. (D)(E) Heatmaps for the TRAPP-LR model and pixel-wise decompositions by LRP for the TRAPP-CNN model for frames 1 and 81. The druggability scores are shown above the heatmaps.

To assess how the TRAPP-LR and TRAPP-CNN models facilitate the identification of more druggable pocket conformations in the TRAPP procedure, an L-RIP simulation was performed using a crystal structure of p38 MAP kinase in the DFG-in conformation (PDB ID: 1r3c)^44^ as the starting structure. The resulting sampled conformations were then analyzed by TRAPP-pocket and druggability assessment (Figure 7C). Through applying perturbations to F169, multiple flips of the Phe in the DFG motif, as well as slight oscillations of the *β*-sheet, were observed in the enhanced sampling MD trajectory. Based on the standardized global descriptors, we observed that the pocket volume, pocket exposure and aromaticity of the starting conformation are close to their corresponding average values calculated from all binding pockets in the NRDLD training set and that they oscillate along the trajectory, whereas the hydrophobicity is higher than average throughout the trajectory. The pocket volume increased around snapshots 20, 40, and 80, which corresponds to the flipping of F169. However, the orientation of D168 does not change in the earlier perturbations (before snapshot 80). The DFG-out conformation is achieved after snapshot 80.

The druggability score of each frame in the L-RIP trajectory was predicted using both the TRAPP-LR and the TRAPP-CNN models. As the druggability scores from the TRAPP-LR and TRAPP-CNN models are squashed into values between 0 and 1, the data points that are not close to the decision boundary would not have significant changes upon conformational change. Here, in order to trace subtle changes in the pocket properties, instead of comparing the druggability score of different conformations, the direct output from the TRAPP-LR and TRAPP-CNN models, before squashing by a logistic or softmax function, is displayed along the trajectory, see Fig 7C. Both models predict the pocket to be druggable and show similar trends in the pre-activated outputs. Thus, the global descriptors and TRAPP-LR are sufficient for tracing druggability in this case. On the other hand, the pocket volume shows the highest correlation to the predicted druggability of the TRAPP-LR and TRAPP-CNN models, indicating the importance of the pocket estimation procedure in TRAPP.

We further computed the RMS distance in terms of the active site residues of each frame to a DFG-out conformation of p38 MAP kinase (1wbs). Most interestingly, the snapshot that had the smallest RMS distance (Frame 81) was the conformation with the highest druggability score. We then investigated the model prediction for frame 1 and 81 through visualization (Figure 7D and E). In snapshot 1, the connection to the sub-pocket is blocked due to the hydrogen bonding between D168 and K53, thus the positive evidence only resides in the main cavity. On the contrary, the DFG motif in snapshot 81 shifts outward and the sub-pocket becomes more open, leading to strong positive evidence in both the main pocket and the cryptic sub-pockets. In addition, extra positive evidence highlights a small hydrophobic cleft near the *β*-sheet, which is connected to the main pocket from above K53 in the pocket estimation procedure, indicating again that the model decision is strongly dependent on the cavity channel extracted from the TRAPP-pocket procedure. In general, we noticed that the druggability of p38 MAP kinase depends strongly on the connection from the main cavity to the cryptic sub-pocket while the orientation of D168 in the flexible loop is not so critical.

Moreover, the evidence for druggable pockets in the DFG-in and DFG-out conformations matches the binding energy hot spots identified using FTMap.^45^ The hotspot regions were discovered by docking 16 different small organic probe molecules and then thresholding the properties of these hotspot clusters to obtain a druggability prediction. A single hot spot at the ATP binding site was detected by mapping unbound structures in the DFG-in loop conformation whereas multiple hot spots, that cover both the ATP binding site and the connected sub-pocket, were identified upon mapping a bound structure in the DFG-out conformation. ^46^ Besides, it has been suggested that cryptic binding sites have a strong binding hot spot in the vicinity and exhibit above-average flexibility around the incipient pockets,^46^ which coincides with the behavior of the cryptic sub-pocket of p38 MAP kinase.

The second example is the cryptic pocket of TEM-1 *β*-lactamase, which is 16 Å away from the center of the active site and only occurs upon binding of a core-disrupting inhibitor. ^47^ The opening of the hydrophobic cryptic pocket between helices 11 and 12 leads to a sequence of linked motions to the catalytic site, which adopts conformations very different from those in the catalytically incompetent conformations. The crystal structures with a closed and an open cryptic pocket (PDB ID: 1jwp and 1pzo) also showed drastic differences in terms of druggability scores (Figure 8A and B). We thus used 1jwp as the starting structure and applied L-RIP perturbations on the L221 of helix 11, which is strongly relocated in the open conformation. The global descriptors and druggability of each conformation obtained from the L-RIP MD trajectory were assessed (Figure 8C). It is apparent from the increase in pocket volume that the cryptic pocket starts to open up after snapshot 40. Additionally, the considerable fluctuations in other physicochemical properties are also a consequence of the small pocket volume as they are normalized to it. The opening of the cryptic pocket correlates with the increase in predicted druggability, especially for the prediction with the TRAPP-CNN model. It is noteworthy that, when comparing the RMS distance between the cryptic site residues of each frame to the open conformation (1pzo), the conformation with the highest similarity (snapshot 68) also has the highest druggability score among all snapshots. By visualizing the model predictions for frames 1 and 68 (Figure 8D and E), a clear difference in terms of pocket volume and hydrogen bonding properties can be identified. The strong positive signal in frame 68 highlights the hydrophobic cryptic pocket, whereas the scattered negative signal regions are located on the protein surface between the two helices.

**Figure 8:**
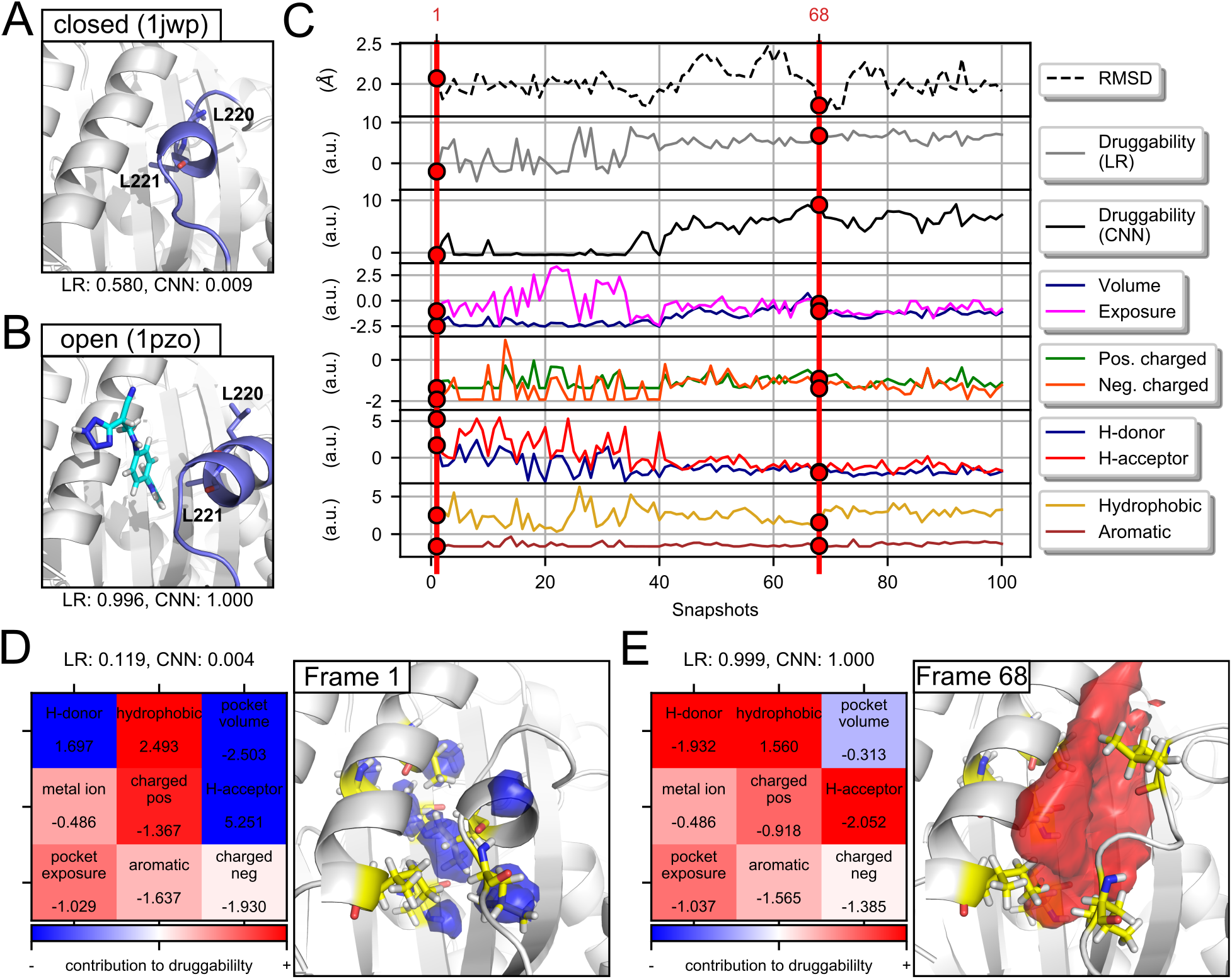
Use of the TRAPP-based models to monitor variations in druggability along an MD simulation of *β*-lactamase. (A)(B) Crystal structures of *β*-lactamase with a closed and an open transient pocket. L220 and L221 are shown in blue, while the bound ligand is shown in cyan. (C) Trace of druggability and other physicochemical properties along an L-RIP trajectory. The RMS distance was calculated between each frame and the open conformation based on the cryptic site residues of *β*-lactamase. The first frame and the frame 68, which has the lowest RMSD to 1pzo, are highlighted. (D)(E) Heatmaps from the TRAPP-LR model and pixel-wise decompositions by LRP for the TRAPP-CNN model for frames 1 and 68. The druggability scores are shown above the heat maps.

## Conclusions

TRAPP is a useful tool for the exploration of binding site conformations and analysis of the flexibility and cavity dynamics. More specifically, the TRAPP-pocket module, which interpolates the atomic information on the binding site onto a grid, enables the identification of conserved and transient pockets upon conformational changes and the comparison of the global properties of the pocket among different conformations. In this work, we have developed two models to predict protein binding pocket druggability and incorporated them into TRAPP to detect fluctuations in druggability upon conformational changes, thus providing a measure for selecting conformations on the basis of druggability. The druggability prediction models are available for use in the TRAPP webserver (https://trapp.h-its.org/).

The two druggability prediction models were built using either the global descriptors or the grid information extracted from TRAPP. The former, TRAPP-LR/SVM provides a linear model trained with logistic regression or a support vector machine using the global descriptors of the binding pockets. The latter, TRAPP-CNN, uses a convolutional neural network to process a grid representation of the properties of the binding pockets.

To train the linear models for druggability prediction based on the global descriptors, namely TRAPP-LR and TRAPP-SVM, the NRDLD dataset^10^ was used. This data set is a consensus data set that has been used for training and validation of state-of-the-art druggability prediction models. To optimize the grid settings, the preprocessing steps, and the hyperparameters for the models together, a pipeline, consisting of preprocessing, standard scaling of the input vectors, and then the LR or SVM model for binary classification, was constructed and a cross-validated grid search was performed. For the final TRAPP-LR and TRAPP-SVM models, which achieve a top mean and a relatively small variance of the cross-validated score in the grid search, a grid edge length of 24 Å, a grid spacing of 0.75 Å, and a hyperparameter C of 1 was used.

The parameters in the trained linear models (Figure 3) indicate the importance of each global descriptor for druggability. The negatively charged and aromatic descriptors are the least important features with small or zero weights in L2 or L1 regularized TRAPP-LR. One of the most intriguing results is that the H-bond donor property has the largest absolute value for its weight, larger than for pocket volume and hydrophobicity, which occupy the second and third places, demonstrating the strong (but negative) contribution of the H-bond donor property to druggability prediction.

TRAPP-CNN, which has detailed grids as inputs and a large amount of adaptive weights, requires a larger training set than TRAPP-LR/SVM. Thus, we augmented the DaPB dataset from the PDBbind refined set 2017, ^25^ in which the positive class is selected by thresholding the properties and the binding affinity of the bound ligand while the negative class was identified using TRAPP-SVM. By projecting the data on to the global descriptors, similar trends were observed as in the NRDLD dataset (Figure 2), indicating that the DaPB dataset is a reasonable surrogate dataset. To overcome overfitting, data augmentations with offline 90° rotation and online random translation were applied to the training data. The network architecture and other hyperparameters were optimized through a cross-validated grid search. The final network contained 3 convolutional layers and 2 fully-connected layers (Figure 4).

The trained models were evaluated on the DaPB test set and the NRDLD test set, both of which are subsets of the dataset that were not used for training. Starting from the performance on the DaPB test set (Table 2), the high specificity was as expected since the parameters of TRAPP-LR are similar to those of TRAPP-SVM, which is responsible for the labeling of the negative class. However, the sensitivity of TRAPP-LR on the DaPB dataset was rather low. This could be due to the fact that the definition of druggable pockets in the NRDLD set is stricter than the definition of druggable (bindable) pockets in the DaPB set. The performance of TRAPP-CNN is close to that of TRAPP-LR, with higher specificity than sensitivity, potentially due to the slightly imbalanced dataset which makes the TRAPP-CNN model favor the negative class. Moreover, all TRAPP-based models have equivalent or superior performance on the NRDLD test set in comparison to the other druggability prediction tools. Both linear models misjudged the same 2 positive and 2 negative data points, whereas the TRAPP-CNN model, which is more precise in detecting the negative class, only missed the 2 positive data points. Thus, the TRAPP-CNN model, although trained on the DaPB bindability dataset, is suitable for druggability prediction.

Apart from the good performance in terms of druggability prediction, a major advantage of our models is the explanatory visualization for model prediction (Figure 5). The contribution of each global descriptor to the druggability prediction from TRAPP-LR is color-coded, which allows easy reasoning of the model output. Furthermore, TRAPP-CNN, which is the first druggability assessment model using the grid representation with detailed spatial information as the input, allows visualization of the regions contributing to the model prediction. We observe that for druggable pockets, TRAPP-CNN mainly identifies extensive hydrophobic patches in the druggable pockets, whereas for less-druggable pockets, it highlights the shallow regions, positively charged groups, or metal ions.

The druggability prediction was integrated as a subroutine in TRAPP-pocket and L-RIP generated trajectories for P38 MAP kinase and *β*-lactamase were analyzed as examples. As the variation of the druggability score is only sensitive when it is close to the decision boundary, we displayed the pre-squashed output from TRAPP-LR and TRAPP-CNN for druggability. The two models show consistent predictions upon conformational changes of the binding pockets. For p38 MAP kinase, the druggability rises as the sub-pocket becomes more connected to the main pocket, which is independent of the position of D168. On the other hand, the striking increase in druggability upon the opening of the cryptic pocket in *β*-lactamase is an ideal example demonstrating the power of combining the TRAPP procedure and druggability prediction.

In summary, we have built three TRAPP-based druggability prediction models, TRAPP-LR, TRAPP-SVN and TRAPP-CNN, which allow the comparison of the druggability of different conformations of a protein binding pocket and the identification of critical sub-regions or properties of a pocket for the corresponding prediction. The druggability assessment is embedded into the pocket estimation procedure in TRAPP, thus providing an efficient platform for selecting protein binding pocket conformations based on druggability scores.

## Supporting information

Labeled DaPB data set in Excel format

Supplementary tables, Table S1 to S7, and figures, Fig.S1, S2, and the description of the seeded region growing (SRG) algorithm

## Acknowledgement

The authors thank Max Horn and Antonia Stank for preliminary work on druggability in TRAPP, and Guido Kanschat and Kai Polsterer for helpful comments. JHY and SH gratefully acknowledge the support of the IWR Scientific Computing-and Molecular Bioscience Master Programs of the Heidelberg University, respectively. This project has received funding from the European Unions Horizon 2020 Framework Programme for Research and Innovation under the Specific Grant Agreement No. 785907 (Human Brain Project SGA2), the EU/EFPIA Innovative Medicines Initiative (IMI) Joint Undertaking K4DD (grant number 115366), and the Klaus Tschira Foundation. This paper reflects only the authors views and neither the IMI nor the European Commission is liable for any use that may be made of the information contained herein.

