## Supplementary tables, Table S1 to S7, and figures, Fig.S1, S2, and the description of the seeded region growing (SRG) algorithm for "Druggability Assessment in TRAPP using Machine Learning Approaches"

**New implementation of the central pocket selection in TRAPP-pocket**

The original central pocket selection procedure is designed to mimic the process of filling up a cavity by traversing from the seed, such as the geometric center of the reference ligand, throughout the whole cavity [1]. However, the traverse directions throughout the grid were predefined in the original implementation, which thus required several traverses with changed order of directions to fully uncover all connected components in the central pocket. To speedup the procedure, seeded region growing (SRG), a method originally designed for image segmentation, is employed.

The essential component of SRG is the use of a list to keep track of the neighboring grid points that satisfy the criterion of homogeneity (in this case, $G(\mathbf{r_i}, p)$ larger than a threshold), enabling the cavity to be captured in a single traverse. The pseudo code of the SRG algorithm is described as follows.

**Algorithm 1:** Seeded Region Growing for Central Pocket Selection

```
Initialize 'OutGrid', same dimension as the input grid. Each element in the input grid
  stores a value G(x̂);
Initialize empty list, 'CheckList';
Set 'OutGrid' at the seed point x_0 to G(x_0);
Put neighbors of the seed point into the 'CheckList';
while the 'CheckList' is not empty do
    Remove first element x̂ from 'CheckList';
    if G(x̂) < δ_thres (criterion of homogeneity) then
        Set 'OutGrid' at x̂ to G(x̂);
        Add unlabeled neighbors of x̂ to 'CheckList';
    else
        Set 'OutGrid' at x̂ to 0;
end
Return 'OutGrid'
```

| **Property** | **Allowed value** |
|---|---|
| Number of non-hydrogen atoms | 10 - 43 |
| Number of H-bond donors | 0 - 6 |
| Number of H-bond acceptors | 0 - 9 |
| Klopman $logP$ | -2.4 - +6.7 |
| Normalized polarity | 0.1 - 0.6 |
| Normalized bond flexibility | 0.1 - 0.4 |
| Binding affinity (in $pK_a$) | 6 - 12 |

Table S1: The list of criteria for selecting drug-like ligands. The normalized polarity is defined as the sum of hydrogen-bond donors and acceptors divided by the number of non-hydrogen atoms, while the normalized bond flexibility is defined as the number of rotatable bonds divided by the total number of non-terminal bonds.

| Ligand ID | Name |
|---|---|
| FE | FE (III) ION |
| FE2 | FE (II) ION |
| FES | FE2/S2 (INORGANIC) CLUSTER |
| MOS | DIOXOTHIOMOLYBDENUM(VI) ION |
| CA | CALCIUM ION |
| MG | MAGNESIUM ION |
| ZN | ZINC ION |
| K | POTASSIUM ION |
| MN | MANGANESE (II) ION |
| NI | NICKEL (II) ION |
| NA | SODIUM ION |
| HEM | PROTOPORPHYRIN IX CONTAINING FE |
| NAP | NADP NICOTINAMIDE-ADENINE-DINUCLEOTIDE PHOSPHATE |
| FAD | FLAVIN-ADENINE DINUCLEOTIDE |
| FMN | FLAVIN MONONUCLEOTIDE |
| NDP | NADPH DIHYDRO-NICOTINAMIDE-ADENINE-DINUCLEOTIDE PHOSPHATE |
| NAD | NICOTINAMIDE-ADENINE-DINUCLEOTIDE |
| VIB | 3-(4-AMINO-2-METHYL-PYRIMIDIN-5-YLMETHYL)-5-(2-HYDROXY-ETHYL)-4-METHYL-THIAZOL-3-IUM |
| PLP | PYRIDOXAL-5'-PHOSPHATE |

Table S2: List of small molecules retained with the protein for the TRAPP-pocket pocket estimation procedure. These metal ions and co-factors are assigned a HETATM type in the PDB format coordinate file.

| PDB ID | Metal ion(s) | PDB ID | Metal ion(s) |
|---|---|---|---|
| 1lox | Fe (+2) | 1xm6 | Zn (+2), Mg (+2) |
| 3etr | Ca (+2), Mo (+6) | 1udt | Zn (+2), Mg (+2) |
| 2cl5 | Mg (+2) | 1r58 | Mn (+2) |
| 1xoz | Zn (+2), Mg (+2) | 1gkc | Zn (+2), Mg (+2) |
| 1r55 | Zn (+2) | 1yqy | Zn (+2) |
| 3f0r | Zn (+2) | 1o5r | Zn (+2) |
| 1oq5 | Zn (+2) | 3pcm | Fe (+3) |
| 1v16 | Mn (+2), K (+1) | 1gpu | Ca (+2) |
| 1wvc | Mg (+2), Ni (+2) | 1qs4 | Mg (+2) |
| 1kc7 | Mg (+2) | 1x9d | Ca (+2) |
| 1px4 | Mg (+2), Na (+1) | 1nnc | Ca (+2) |
| 1ec9 | Mg (+2) | 1icj | Ni (+2) |
| 1e9x | Fe (+3) | 1hqg | Mn (+2) |
| 1sqi | Fe (+3) | 2gsu | Zn (+2) |
| 1r9o | Fe (+3) | 2gyi | Mg (+2) |
| 1kvo | Ca (+2) | - | - |

Table S3: List of PDB files in the NRDLD dataset containing proteins with metal ions in the binding pocket. The PDB ID, metal ions, and their corresponding charges (in e units) are shown.

| **Property** | **Definition** |
| --- | --- |
| Pocket volume | $\sum_{i=1}^{N}(l_p)^3 \times [G(\mathbf{r_i}, p) > 0]$ |
| Protein-exposed surface area | $\sum_{i=1}^{N}(l_p)^2 \times [G(\mathbf{r_i}, p) = 0 \wedge G(\mathbf{r_{i_{(-1)}}}, p) > 0]$ |
| Solvent-exposed surface area | $\sum_{i=1}^{N}(l_p)^2 \times [G(\mathbf{r_i}, p) = 0 \wedge G(\mathbf{r_{i_{(-1)}}}, p) > -1]$ |
| Pocket exposure | $\frac{\text{Solvent-exposed surface area}}{\text{Protein-exposed surface area}} \times 100(\%)$ |
| Positively charged | $\sum_{i \in \mathcal{Q}}(l_p)^3 \times G^{ch}(\mathbf{r_i}, p) \times [G(\mathbf{r_i}, p) > 0]$ |
| Negatively charged | |
| Hydrogen-bond donor | |
| Hydrogen-bond acceptor | $\sum_{i=1}^{N}(l_p)^3 \times G^{at}(\mathbf{r_i}, p) \times [G(\mathbf{r_i}, p) > 0]$ |
| Hydrophobic | |
| Aromatic | |
| Metal ion | |

Table S4: Definitions of the global descriptors generated in the TRAPP-pocket procedure.The grid contains $N$ grid points in one channel. A grid point is denoted as $\mathbf{r_i}$, where $i = 1 \cdots N$. $p$ represents a particular protein structure. $G(\cdot)$, $G^{ch}(\cdot)$, and $G^{at}(\cdot)$ are the distribution functions for cavity, charged atoms and other atomic properties, respectively. The grid spacing is denoted as $l_p$, thus a unit volume and a unit surface area in the grid are $(l_p)^3$ and $(l_p)^2$. $\mathbf{r_{i_{(-1)}}}$ denotes the grid point that is examined before the current grid point $\mathbf{r_i}$ in the region growing algorithm. The indicator function $[\cdot]$ represents a function that outputs 1 if the condition is satisfied, and 0 otherwise. The set $\mathcal{Q}$ holds all grid points that are within the pocket.

| **Hyperparameter** | **values/options** |
|---|---|
| Grid spacing Å | $0.5, 0.75, 1.0$ |
| Grid edge length Å | $21, 24, 27$ |
| Vol. norm. | yes/no |
| Skip metal | yes/no |
| C | 0.01, 0.1, 1, 10, 100 |

Table S5: Hyperparameters tuned in the TRAPP-SVM and TRAPP-LR pipelines. Vol. norm.: normalization of the physicochemical properties to the pocket volume. Skip metal: removal of the metal ion property from the input features.

The hyperparameter tuning was performed in consecutive runs of cross-validated grid search instead of one grid search for all combinations of hyperparameters, due to the combinatorial increase in run time. For the training of TRAPP-LR and TRAPP-SVM, the $F_1$ score was used to compare the performance between models with varied hyperparameters. We performed the first round of the grid search over 60 similar architectures with varied depth and width of the network as shown in Table S6. In the second round of the grid search, the learning rate and weight decay for regularization were optimized as in Table S7.

| Conv layers \ Extra FC layer | 0 | 16 | 256 |
|---|---|---|---|
| (16, 16, 32, 0, 0) | 0.767 (±0.042) | **0.784 (±0.011)** | 0.502 (±0.717) |
| (16, 32, 32, 0, 0) | 0.779 (±0.034) | **0.782 (±0.052)** | **0.785 (±0.050)** |
| (16, 32, 64, 0, 0) | 0.775 (±0.014) | **0.782 (±0.026)** | 0.521 (±0.738) |
| (32, 32, 64, 0, 0) | 0.519 (±0.734) | 0.769 (±0.069) | **0.792 (±0.055)** |
| (32, 64, 64, 0, 0) | 0.525 (±0.742) | 0.521 (±0.737) | 0.522 (±0.738) |
| (32, 64, 128, 0, 0) | 0.748 (±0.042) | **0.794 (±0.024)** | 0.775 (±0.027) |
| (16, 16, 32, 32, 0) | 0.774 (±0.026) | 0.472 (±0.672) | **0.799 (±0.045)** |
| (16, 16, 32, 64, 0) | 0.768 (±0.015) | 0.774 (±0.059) | 0.768 (±0.056) |
| (16, 32, 32, 64, 0) | 0.755 (±0.039) | 0.526 (±0.744) | 0.265 (±0.749) |
| (16, 32, 64, 64, 0) | 0.764 (±0.043) | 0.502 (±0.710) | 0.495 (±0.701) |
| (32, 32, 64, 64, 0) | **0.787 (±0.031)** | 0.527 (±0.746) | 0.774 (±0.023) |
| (32, 32, 64, 128, 0) | 0.518 (±0.733) | 0.515 (±0.728) | 0.534 (±0.755) |
| (32, 64, 64, 128, 0) | 0.509 (±0.723) | 0.245 (±0.691) | 0.527 (±0.746) |
| (32, 64, 128, 128, 0) | 0.779 (±0.050) | 0.762 (±0.055) | 0.238 (±0.673) |
| (16, 16, 32, 32, 64) | 0.757 (±0.044) | **0.787 (±0.036)** | 0.263 (±0.745) |
| (16, 16, 32, 64, 64) | 0.763 (±0.068) | 0.762 (±0.050) | **0.781 (±0.006)** |
| (16, 32, 32, 64, 64) | 0.754 (±0.049) | 0.507 (±0.718) | 0.523 (±0.740) |
| (32, 32, 64, 64, 128) | 0.511 (±0.724) | 0.778 (±0.033) | 0.538 (±0.761) |
| (32, 32, 64, 128, 128) | 0.774 (±0.062) | 0.249 (±0.704) | 0.249 (±0.704) |
| (32, 64, 64, 128, 128) | 0.745 (±0.096) | 0.484 (±0.686) | 0.772 (±0.045) |

Table S6: 3-fold cross-validated grid search for optimizing the architecture of TRAPP-CNN. The five element tuple in each row indicates the number of convolutional filters used in each layer, where zero indicates the layer does not exist. In total, there are 20 configurations for the convolutional layers. The three columns correspond to the configuration of the extra fully-connected layer before the output layer, where 0 indicates no extra fully-connected layer, and 16 and 256 are the number of hidden nodes in this fully-connected layer. The mean of the 3-fold cross validation F1 score is shown together with the standard deviation in brackets. The performance of the top configurations is marked in bold.

| Weight decay \ Learning rate | 1.00E-04 | 1.00E-03 | 1.00E-02 |
|---|---|---|---|
| 1.00E-05 | 0.793 (±0.060) | 0.256 (±0.724) | 0.000 (±0.000) |
| 1.00E-04 | 0.789 (±0.044) | **0.802 (±0.050)** | 0.000 (±0.000) |
| 1.00E-03 | 0.796 (±0.047) | 0.773 (±0.058) | 0.000 (±0.000) |

Table S7: 3-fold cross-validated grid search for optimizing the learning rate and weight decay for TRAPP-CNN. The network architecture is shown in Figure 4. The mean of the 3-fold cross validation F1 score is shown together with the standard deviation in brackets. The performance of the top configurations is marked in bold. All of the $F1$ scores obtained when using a learning rate = 0.01 are ill-defined and thus shown as 0.000

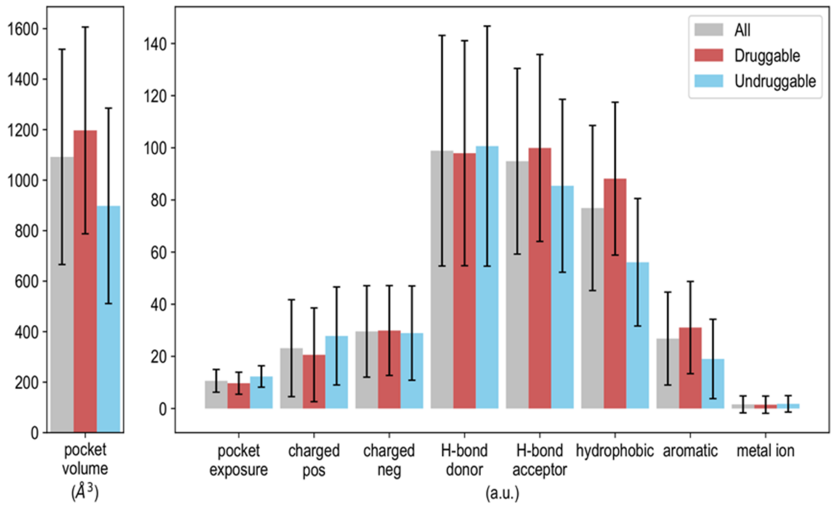


Figure S1: Visualization of the NRDLD[2] dataset : Mean and standard deviation of each global property computed using all, druggable and less-druggable protein structures in the dataset.

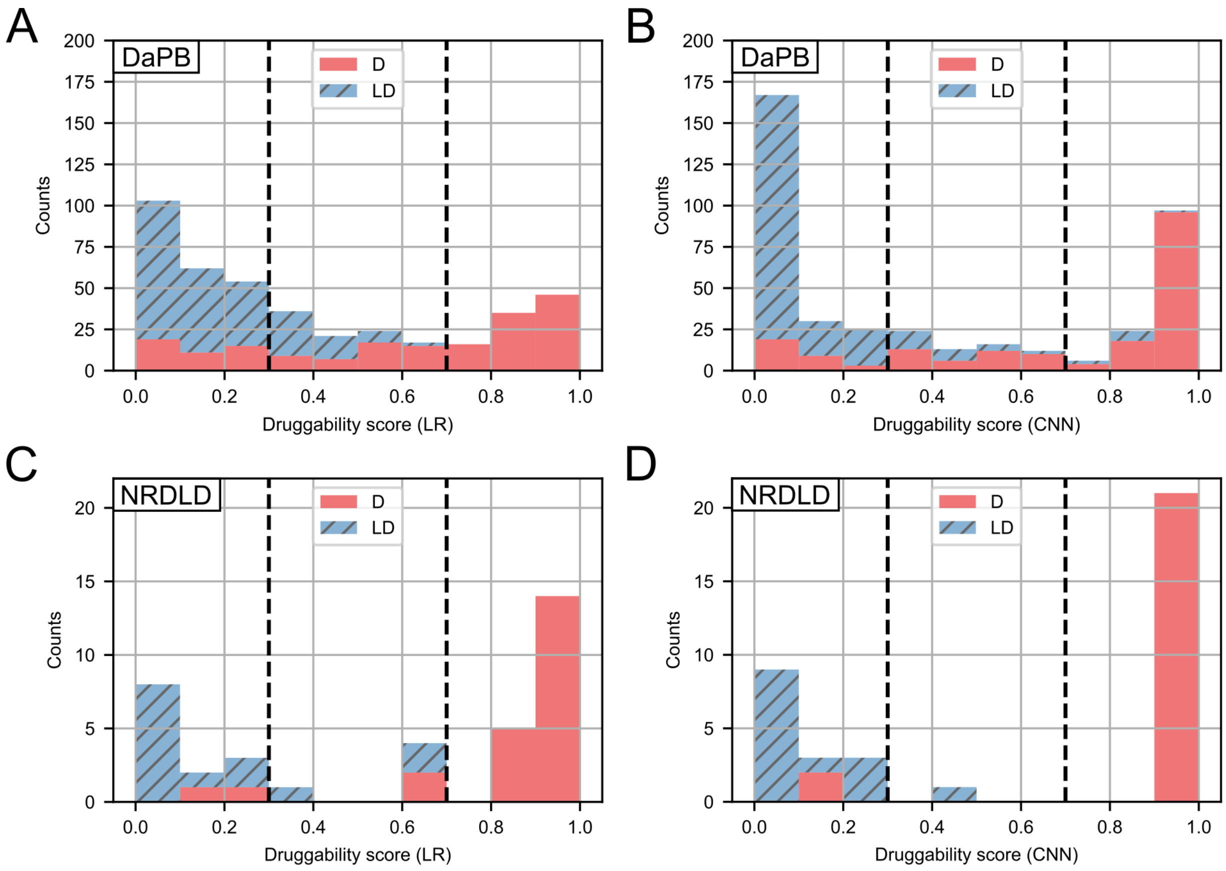


Figure S2: Distribution of the druggability scores predicted by TRAPP-LR(A,C) and TRAPP-CNN(B,D) on the DaPB(A,B) and NRDLD(C,D) test sets. The druggable and less-druggable pockets are represented in red filled and blue hatched bars. In the region between the dashed lines at druggability scores of 0.3 and 0.7 the predictions are uncertain.
